# The *Iroquois* (*Iro/Irx*) homeobox genes are conserved Hox targets involved in motor neuron development

**DOI:** 10.1101/2024.05.30.596714

**Authors:** Catarina Catela, Stavroula Assimacopoulos, Yihan Chen, Konstantinos Tsioras, Weidong Feng, Paschalis Kratsios

## Abstract

The *Iroquois (Iro/Irx)* homeobox genes encode transcription factors with fundamental roles in animal development. Despite their link to various congenital conditions in humans, our understanding of *Iro/Irx* gene expression, function, and regulation remains incomplete. Here, we conducted a systematic expression analysis of all six mouse *Irx* genes in the embryonic spinal cord. We found five *Irx* genes (*Irx1, Irx2, Irx3, Irx5,* and *Irx6*) to be confined mostly to ventral spinal domains, offering new molecular markers for specific groups of post-mitotic motor neurons (MNs). Further, we engineered *Irx2, Irx5,* and *Irx6* mouse mutants and uncovered essential but distinct roles for *Irx2* and *Irx6* in MN development. Last, we found that the highly conserved regulators of MN development across species, the HOX proteins, directly control *Irx* gene expression both in mouse and *C. elegans* MNs, critically expanding the repertoire of HOX target genes in the developing nervous system. Altogether, our study provides important insights into *Iro/Irx* expression and function in the developing spinal cord, and uncovers an ancient gene regulatory relationship between HOX and *Iro/Irx* genes.

## Introduction

The *Iroquois homeobox (Irx)* genes encode a family of conserved homeodomain transcription factors^1^. They play crucial roles in the regulation of various developmental processes, particularly in cell specification and differentiation during embryonic development^2^. First discovered in the fruit fly *Drosophila melanogaster,* the *Irx* genes were named after their distinct mutant phenotype; *Iroquois* mutant flies display a wide band of sensory organs across the central region of mesothorax – a pattern resembling the hairstyle worn by members of the Iroquois Confederacy^3^. With the exception of the nematode *Caenorhabditis elegans*, which has a single ancestral *Iroquois* gene *(irx-1)*, *Irx* genes are typically organized in genomic clusters. Three genes (*ara, caup, mirr*) are present in a single cluster in *Drosophila*, whereas six genes in mice (*Irx1-6*) and humans (*IRX1-6*) are found in two separate chromosomal clusters (**Fig. 1A-B**). Irx proteins bind DNA through a highly conserved 63 amino acid-long homeodomain of the TALE (three-amino acid loop extension) class and carry a distinct 13 amino acid domain (Iro box) involved in protein-protein interactions (**Fig. 1C**) ^4^.

**Figure 1.**
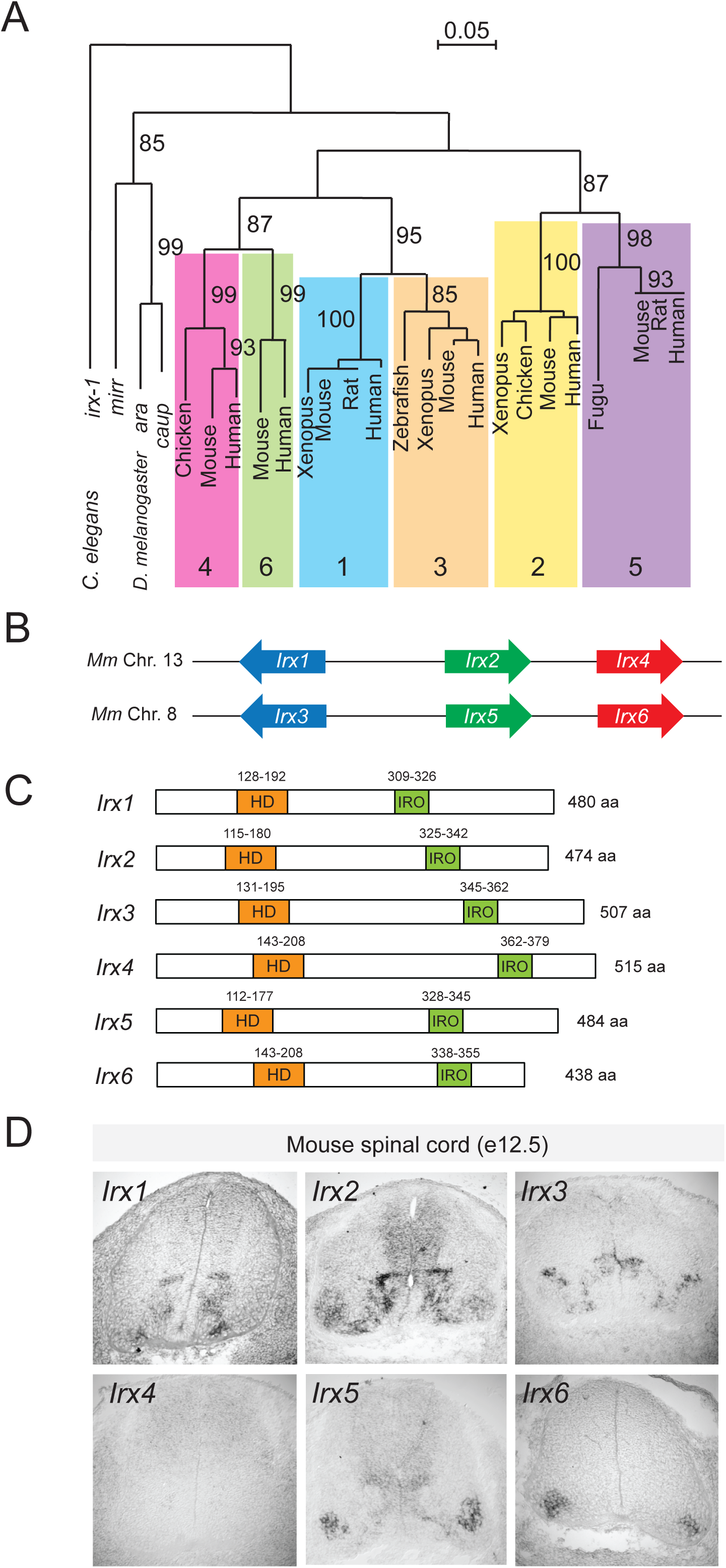
Expression pattern of all six *Irx* genes in mouse spinal motor neurons. (A) Phylogenetic tree of the *Iro/Irx* gene family. Adapted from ^101^. Scale bar: percentage of sequence divergence. Numbers above branches: percentage of times the branch was found in 200 bootstrap replicates. Brackets highlight the clustering of vertebrate genes into six orthologous groups. (B) Schematic of the two *Irx* chromosomal clusters in mice. (C) Schematic of the mouse Irx protein domains. Each protein has a highly conserved 63 amino acid-long homeodomain and a distinct 16 amino acid-long Iro box domain. (D) RNA ISH analysis of *Irx1*, *Irx2*, *Irx3*, *Irx4*, *Irx5*, and *Irx6* on cross-sections of e12.5 wild-type mouse spinal cords. Brachial (C4-T1) domain. N = 4.

Emerging evidence shows that *Irx* gene expression spans large territories in early embryos, but becomes restricted to distinct cell types and tissues at later stages of development^1,2^. As such, *Irx* genes not only control embryonic patterning and cell specification, but also play later roles in tissue differentiation and function. In *Drosophila*, *Irx* genes specify large territories in the developing eye and wing disc^1,5,6^. In vertebrates, they control cell specification and function in the developing heart^7^, lung ^8^, kidney ^9,10^, pancreas^11^, gonad^12^ and limb^13^. Mutations in human *IRX* genes can lead to a range of phenotypic effects, such as heart defects, craniofacial abnormalities, and cancer ^7,14–16^. Much of our current knowledge on *Irx* genes, however, derives from studies in the murine heart, where a systematic expression pattern analysis of all six *Irx* genes has been complemented with individual and compound *Irx* mutant analyses ^7,17,18^. These studies revealed both essential and redundant functions for *Irx* genes, and further demonstrated that Irx proteins can either act as activators or repressors of gene expression. Beyond the developing heart, systematic expression and mutant analyses for *Irx* genes are currently lacking in other vertebrate tissues (e.g., kidney, brain, spinal cord), limiting our understanding of how these highly conserved transcription factors control animal development.

In the nervous system, *Iro/Irx* genes play crucial roles during early and late stages of development ^2^. For example, growing evidence suggests their requirement for the initial specification of vertebrate neuroectoderm, the first step of nervous system development. In *Xenopus*, *Iro* genes specify the neural territory during gastrulation ^19–21^. Both in *Drosophila* and vertebrates, *Iro/Irx* genes are known to regulate proneural genes essential for the development of neuroectodermal progenitor cells ^19,22,23^. Subsequently, *Iro/Irx* genes subdivide the developing neuroectoderm and promote regional identities along the dorsoventral (D-V) axis of the developing spinal cord and anterior-posterior axis of the brain^2,24–26^. They do so by generating sharp boundaries between dividing progenitor cells through mutual cross-repression with other transcription factors. In progenitor cells, Irx3 cross-represses Olig2 in the spinal cord, Six3 in the forebrain, and Hnf1 in the hindbrain ^27–30^.

During later stages of neural development, *Iro/Irx* genes are best characterized in the developing retina, where *Drosophila* and mouse studies revealed essential roles in neurogenesis ^31,32^ and cell specification ^1,6,33,34^. In the mouse retina, *Irx4* controls axon guidance, whereas *Irx5* and *Irx6* are necessary for interneuron terminal differentiation^35–37^.

Moreover, *Irx* genes have been implicated in cerebellum formation^38^, serotonergic neuron differentiation^39^, and postnatal neurogenesis in the hypothalamus^40^. Human *IRX3* and *IRX5* have been associated with obesity, and mouse studies demonstrated that hypothalamic *Irx3* and *Irx5* are involved in feeding behavior and metabolism ^41–44^. Further, *IRX5* mutations cause a recessive congenital disorder affecting brain, face, blood, and heart development^14^. Altogether, multiple studies to date have reported essential roles for *Irx* genes in various parts (retina, cerebellum, hypothalamus) of the developing central nervous system (CNS), except the spinal cord, where our knowledge of *Irx* gene function remains rudimentary.

How is *Iro/Irx* gene expression controlled during development? During early patterning, studies in *Drosophila* showed that *Iro* genes are activated by multiple signaling pathways, such as Hedgehog, Wingless, JAK/STAT, and EGFR ^45,46^. In the vertebrate neural plate, *Irx/Iro* gene expression is activated by Wnt and repressed by BMP signaling^21,47^. During later stages of CNS development, *Irx2* is a target of the FGF8/MAP kinase cascade in the midbrain and *Irx3* is repressed by Shh signaling in the spinal cord^38,48^. However, how *Irx/Iro* gene expression is regulated in the context of post-mitotic neurons remains largely unknown.

Here, we report a detailed expression map of all six mouse *Irx* genes in the embryonic spinal cord, revealing unique and overlapping expression patterns in distinct populations of post-mitotic neurons. Through CRISPR/Cas9 gene editing, we engineered mouse mutants for *Irx2, Irx5,* and *Irx6*, and uncover essential (*Irx2*, *Irx6)* and dispensable *(Irx5)* roles in spinal MN development. Lastly, we provide evidence – both in *C. elegans* and mice – that Hox transcription factors act directly to control *Irx* gene expression in post-mitotic MNs, exposing an ancient gene regulatory relationship between two highly conserved families of clustered transcription factors.

## RESULTS

### A detailed map of *Irx* gene expression in mouse spinal motor neurons

In mice, *Irx* genes are expressed in various domains of the developing nervous system ^49–53^, but their expression in post-mitotic neurons is poorly defined. Here, we systematically investigated the expression of all six mouse *Irx* genes in the developing spinal cord at the brachial (also referred to as “cervical”) domain (C4-T1) (**Fig. 2A**). We focused on embryonic day 12.5 (e12.5), as the majority of post-mitotic spinal neurons have been generated by that stage. RNA *in situ* hybridization (ISH) showed that *Irx1, Irx2, Irx3* and *Irx5* are mostly confined to medial and ventral regions of the spinal cord, whereas *Irx6* is specific to the ventrolateral region (**Fig. 1D**). Consistent with a previous report at stages earlier than e12.5^49^, we did not detect expression of *Irx4* (**Fig. 1D**), the most divergent member of the vertebrate *Irx* gene family^54^.

**Figure 2.**
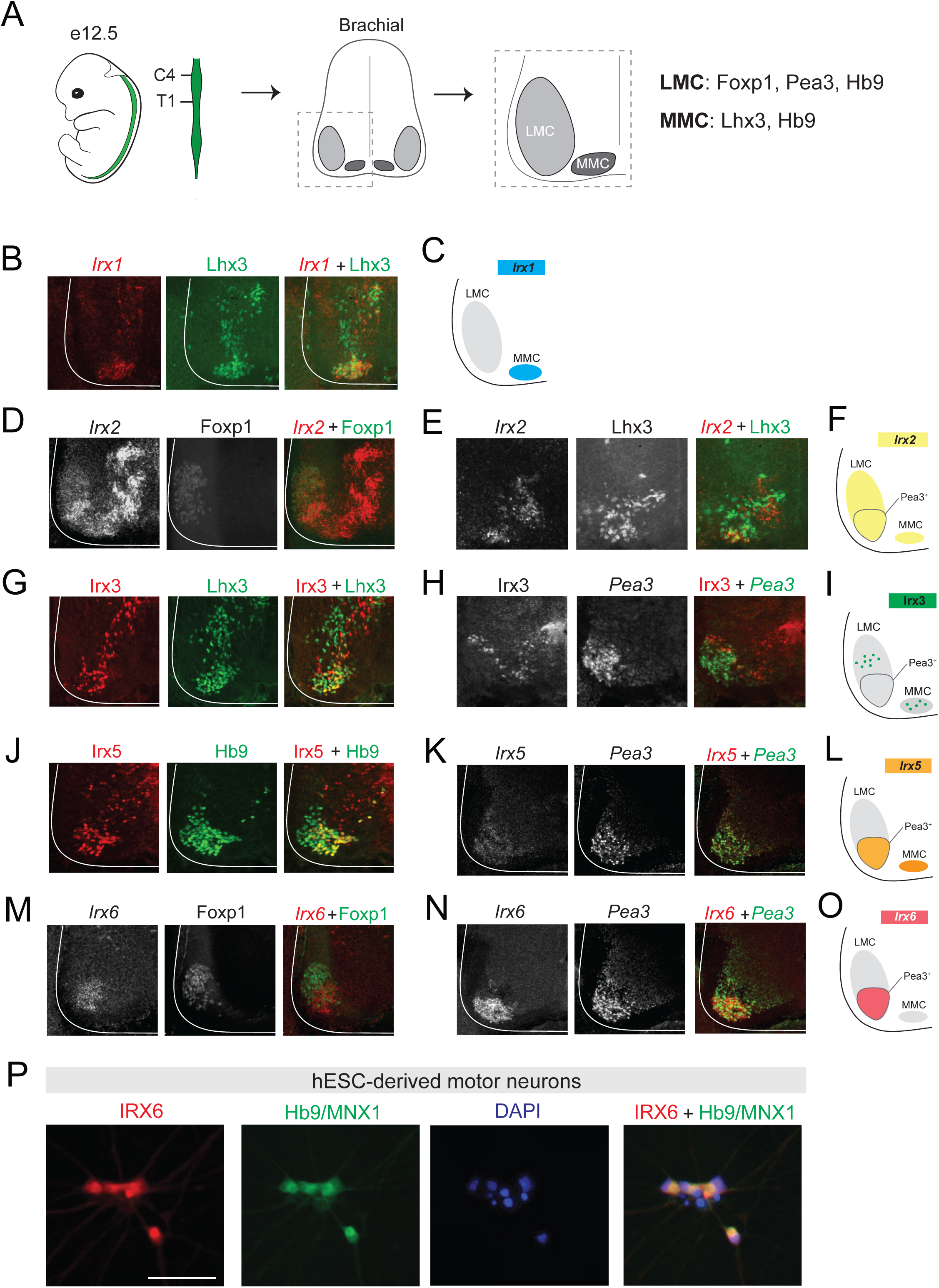
Expression of *Irx1, Irx2, Irx3, Irx5,* and *Irx6* in specific motor columns. (A) Brachial spinal domain (C4-T1) in an e12.5 mouse. Cross-section: LMC and MMC neurons at the ventral region of the spinal cord. (B) *Irx1* FISH coupled with Lhx3 antibody staining shows *Irx1* expression (red) is detected in Lhx3-expressing MNs (green) of the MMC. (C) Schematic of *Irx1* expression. (D-E) *Irx2* FISH combined with immunostaining for Foxp1 (D) or Lhx3 (E) reveals *Irx2* expression (red, also converted to white for better contrast) is detected in LMC (D) and MMC (E) neurons. (F) Schematic of *Irx2* expression. (G) Double immunostaining for Irx3 and Lhx3 shows Irx3 (red) is expressed in a small population of Lhx3-positive MNs (green). (H) Irx3 antibody staining combined with *Pea3* RNA FISH reveals almost no co-localization of Irx3 protein (red) to *Pea3*-expressing MNs (green) of the ventral LMC but expression in a small population of dorsal LMC MNs. (I) Schematic of *Irx3* expression. (J) Double immunofluorescence reveals Irx5 expression (red) in Hb9-positive cells (green). (K) Two-color RNA FISH shows *Irx5* expression (red) is detected in *Pea3*-expressing MNs of the ventral LMC. (L) Schematic of *Irx5* expression. (M) *Irx6* RNA (red) colocalizes with Foxp1 protein (green) in the ventral LMC. (N) Two-color RNA FISH reveals *Irx6* expression (red) in *Pea3*-expressing MNs. (O) Schematic of *Irx3* expression. (P) Representative images of HUES3 HB9::GFP-derived MNs upon immunostaining for HB9/MNX1 and IRX6 on day 21 of differentiation. Scale bar, 50 μm.

As the majority of *Irx* expression occurs in the ventral region of the spinal cord, which is known to contain post-mitotic motor neurons (MNs), we focused our analysis on these cells. At the brachial domain (C4-T1) of the spinal cord, MNs are mainly organized into two columns. The lateral motor column [LMC] contains forelimb-innervating MNs necessary for reaching, grasping, and locomotion, whereas the medial motor column [MMC] contains axial muscle-innervating MNs required for postural control (**Fig. 2A**) ^55^. At e12.5, post-mitotic MNs (LMC or MMC) can be unambiguously distinguished by suing established molecular markers for motor columns (LMC: Foxp1; MMC: Lhx3; LMC and MMC: Hb9; a subpopulation of LMC: Pea3) ^56–58^ (**Fig. 2A**). To generate a detailed map of *Irx* expression in post-mitotic MNs of the e12.5 brachial spinal cord, we evaluated expression of five mouse *Irx* genes (*Irx1, Irx2, Irx3, Irx5. Irx6*) with RNA fluorescent ISH (FISH).

By coupling RNA FISH for *Irx1* with immunofluorescence for the MMC marker Lhx3, we found that *Irx1* is expressed in MMC neurons (**Fig. 2B-C**). Through similar double labeling strategies that rely on RNA FISH and/or immunofluorescence, we determined that: (a) *Irx2* is expressed in MMC and LMC neurons (including the Pea3 pool of MNs) (**Fig. 2D-F, Supplementary Fig. 1A**), (b) *Irx3* mRNA and protein are present in MMC neurons and in a small population of LMC neurons (located dorsally to the Pea3 MN pool) (**Fig. 2G-I**, **Supplementary Fig. 1B**), (c) *Irx5* mRNA and protein are detected in MMC neurons and the Pea3 pool (**Fig. 2J-L**, **Supplementary Fig. 1C**), and (d) *Irx6* mRNA is remarkably specific to the Pea3 pool within the LMC neurons (**Fig. 2M-O**). In addition to MNs, *Irx1, Irx2, Irx3*, and *Irx5* are expressed in other spinal cell types close to the midline, likely progenitor cells and various types of interneurons (**Fig. 2**). Altogether, this analysis provides a detailed expression map of *Irx* genes in post-mitotic spinal MNs, offering new molecular markers for embryonic MMC and LMC neurons (**Fig. 2**).

At the protein level, we evaluated nine commercially available Irx antibodies by performing immunofluorescence staining in the mouse spinal cord (**Supplementary Table 1**). We found two antibodies that display specificity; one against Irx3 and another against Irx5 (**Supplementary Table 1**). Antibody staining against Irx3 and Irx5 yielded expression patterns similar to the ones observed with *Irx3* and *Irx5* RNA ISH (**Fig. 1C, 2G, 2J**).

### IRX6 is expressed in human motor neurons derived from embryonic stem cells

To determine whether IRX expression in motor neurons is a conserved feature from mice to human, we generated MNs from a human embryonic stem cell (hESC) line (HUES3 HB9::GFP) ^59^. Following differentiation of the HUES3 line and immunofluorescence (see Materials and Methods), we found that IRX6 co-localizes with HB9 in human post-mitotic MNs (day 21 in culture) (**Fig. 2P**). Because the differentiation protocol we used is known to generate LMC neurons ^60^, IRX6 is likely expressed in human LMC neurons, consistent with the *Irx6* expression in the mouse spinal cord (**Fig. 2M-O**). In addition to detecting IRX6, we also tested several commercially available antibodies against human IRX2, IRX3, and IRX5, but failed to detect specific signal in hESC-derived MNs (**Supplementary Table 1**).

### *Irx2, Irx5* and *Irx6* are expressed in LMC neurons at brachial and lumbar domains

To determine *Irx* expression in MNs along the rostrocaudal axis of the spinal cord, we examined thoracic and lumbar domains at e12.5. Similar to brachial domains, MMC neurons of thoracic and lumbar domains also express *Irx2* (**Supplementary Fig. 2**). Hence, *Irx2* marks post-mitotic MMC neurons along the embryonic spinal cord. Unlike MMC neurons, LMC neurons are only present at brachial and lumbar domains, from which they innervate forelimb and hindlimb muscles, respectively^61^. However, the extent of molecular similarity between brachial and lumbar MNs remains poorly defined as most studies focus on brachial MNs^62^. We found that *Irx2, Irx5,* and *Irx6* are not only expressed in brachial LMC neurons, but also in their lumbar counterparts (**Supplementary Fig. 2**), thereby constituting shared LMC markers expressed by both brachial and lumbar LMC neurons.

### *Irx2* is required for limb-innervating motor neuron development

Unlike *Irx3, Irx5,* and *Irx6*, *Irx2* is expressed in most, if not all, LMC neurons in the brachial domain of the spinal cord (**Fig. 2D-F**). To test whether *Irx2* is necessary for LMC neuron development, we used CRISPR/Cas9 to generate *Irx2* mutant mice, herein referred to as *Irx2 ^15bp/15bp^* because they carry a 5bp-long deletion in the 2^nd^ exon of *Irx2* (**Fig. 3A**) (see Materials and Methods). Consistent with previously described *Irx2* KO mice generated by classic Cre/loxP recombination ^63^, homozygous *Irx2 ^15bp/15bp^* are viable and fertile. The 5bp deletion is predicted to cause a frameshift in the open reading frame (at codon 122), introducing a premature termination codon (**Fig. 3A-B**). Such a frameshift drastically changes the amino acid composition of the Irx2 homeodomain (**Fig. 3B, Supplementary File 1**), which is essential for DNA-binding.

**Figure 3.**
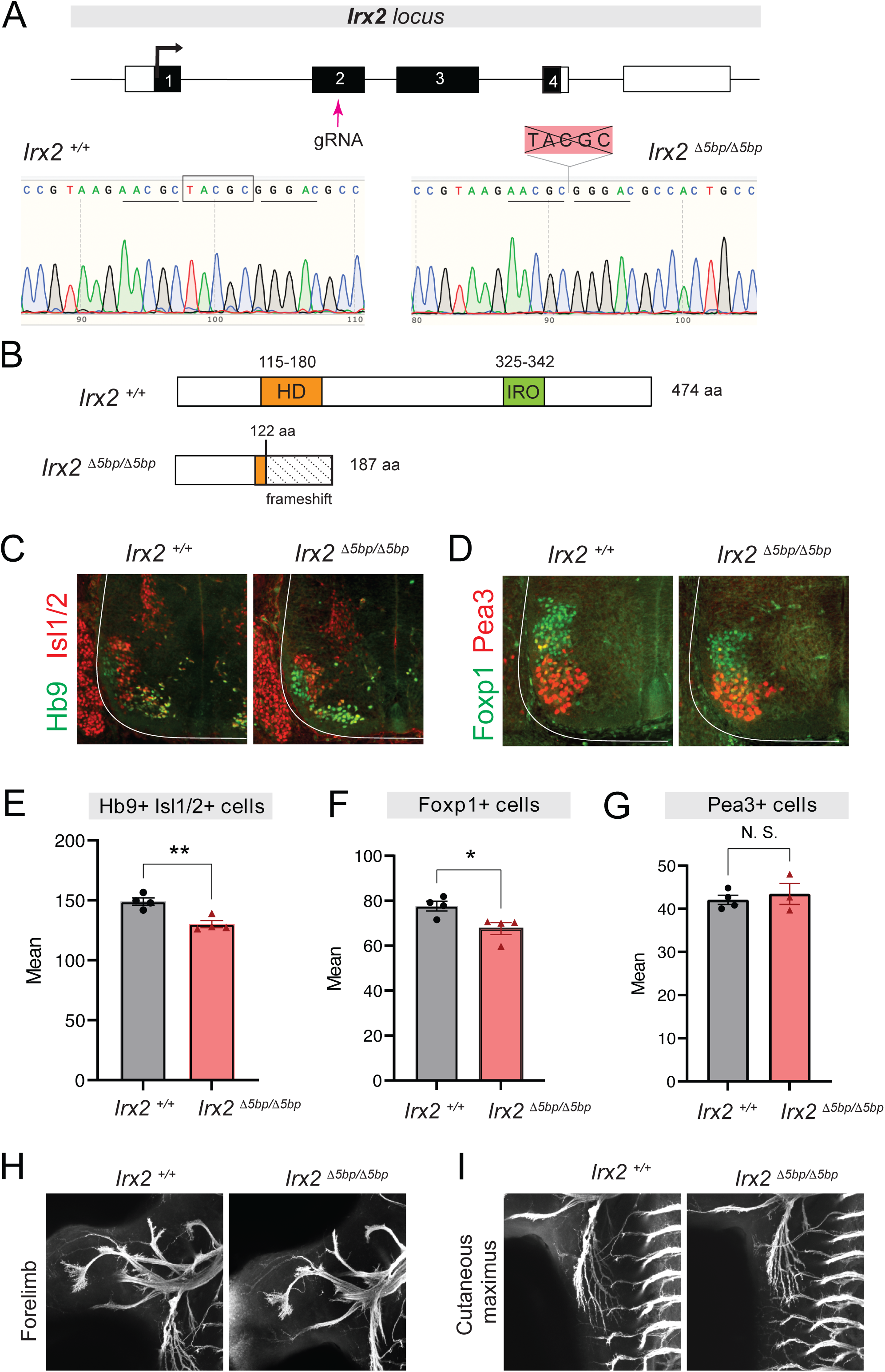
Generation and characterization of *Irx2* mutant mice. (A) A gRNA targets exon 2 of *Irx2*. Representative chromatograms of genotyped mice demonstrate the 5bp (TACGC) deletion in *Irx2* ^Δ*5bp/*Δ*5bp*^ mice. (B) Wild-type and truncated Irx2 proteins in the *Irx2* ^Δ*5bp*^ mouse. See text for details. (C-D) Double immunostaining for Hb9 (green) and Isl1/2 (red) (panel C) or Foxp1 (green) and Pea3 (red) (panel D) in e12.5 wild-type and *Irx2* ^Δ*5bp/*Δ*5bp*^ spinal cords. (N=4). (E-G) Quantification of Hb9+/Isl1/2+, Foxp1+, or Pea3+ MNs in *Irx2* ^Δ*5bp/*Δ*5bp*^ mice at e12.5 (N=4). * p < 0.05, ** p < 0.01, *** p < 0.001, N. S.: Not significant. (H-I) MN axons labeled with Hb9::GFP in control and *Irx2* ^Δ*5bp/*Δ*5bp*^ mice at e12.5 (N=4). Projections to forelimb (H) and cutaneous maximus (I) muscle.

In the brachial domain of *Irx2 ^15bp/15bp^* spinal cords at e12.5, we found a significant reduction in the total number of MNs, assessed by the co-expression of Hb9 and Isl1/2 (**Fig. 3C, E**). Using the LMC-specific marker FoxP1, we found that this reduction specifically affects LMC neurons (**Fig. 3D, F**). To identify which subpopulation of LMC neurons is affected, we stained for Pea3 (ETV4), which marks a specific subgroup of LMC neurons that innervate back muscles in mice^57^. We found no difference in the number of Pea3-expressing MNs between control (*Irx2^+/+^*) and *Irx2 ^15bp/15bp^* mice (**Fig. 3G**). We conclude that the number of LMC neurons located outside the Pea3 pool (Foxp1+ Pea3-) is reduced in the brachial spinal cord of *Irx2* mutant animals.

### Axonal projections and forelimb grip strength are unaffected in *Irx2* mutants

Because *Irx* genes are involved in axon guidance in the retina^36^, we wondered whether *Irx2* is required for motor axon guidance. To visualize MN axons, we crossed the *Irx2 ^15bp/15bp^* animals to *Hb9(Mnx1)::GFP* mice and performed wholemount GFP immunostaining at e12.5. Compared to control mice, MN axonal projections to forelimb and trunk (cutaneous maximus) muscles appear grossly normal in *Irx2 ^15bp/15bp^* animals (**Fig. 3H-I**). Next, we reasoned that the reduced numbers of LMC neurons in *Irx2 ^15bp/15bp^* mice may affect forelimb grip strength, as LMC neurons specifically innervate forelimb muscles. However, we did not observe a statistically significant effect in adult *Irx2 ^15bp/15bp^* animals compared to controls (**Supplementary Fig. 3**). Altogether, we observe a modest decrease in the number of brachial LMC neurons in *Irx2 ^15bp/15bp^* mice, but no significant effects on axon guidance and forelimb grip strength.

### *Irx5* is not required for brachial motor neuron generation and axon guidance

Like *Irx2*, *Irx5* is expressed in LMC neurons (**Fig. 2J-L**). To test whether *Irx5* controls aspects of LMC development, we employed CRISPR/Cas9 to generate *Irx5* mutant mice, which carry a 5bp-long deletion in the 1^st^ exon of *Irx5* (**Fig. 4A-B**) (see Materials and Methods). Homozygous *Irx5 ^15bp^*^/^*^15bp^* mice are viable. This deletion is predicted to result in a premature termination codon, likely generating a truncated protein that lacks both the HD and IRO domains (**Fig. 4B, Supplementary File 1**). Antibody staining showed dramatically reduced Irx5 protein expression in *Irx5^15bp/15bp^* spinal cords at e12.5 (**Fig. 4C**), indicating *Irx5^15bp^*is a loss-of-function allele. However, quantification of either all brachial MNs (Hb9+ cells) (**Fig. 4D, F**), only LMC neurons (Foxp1+ cells) (**Fig. 4E, G**), or MNs of the Pea3 population (Pea3+ cells) (**Fig. 4E, H**) did not reveal any differences between control and *Irx5 ^15bp^*^/^*^15bp^* animals at e12.5. Further, MN axonal projections to forelimb and trunk (cutaneous maximus) muscles appear grossly normal in *Irx5 ^15bp^*^/^*^15bp^* animals (**Fig. 4I-J**). We conclude that *Irx5* is not required for brachial MN generation or axon guidance, raising the possibility that *Irx5* acts redundantly with *Irx6* given their overlapping expression in LMC neurons (**Fig. 2**).

**Figure 4.**
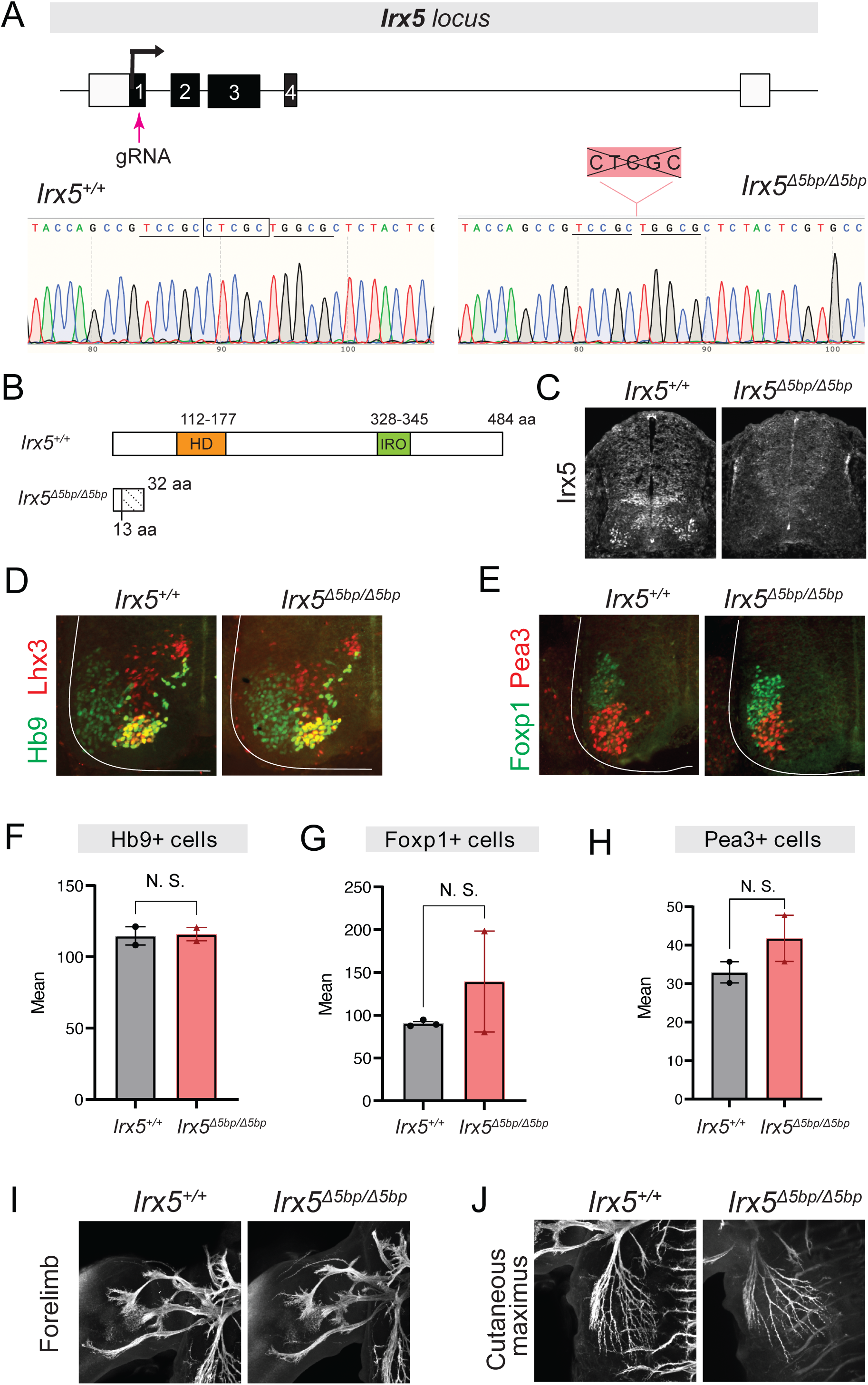
Generation and characterization of *Irx5* mutant mice. (A) A gRNA targets exon 1 of *Irx5*. Representative chromatograms of genotyped mice illustrate the 5 bp (CTCGC) deletion in *Irx5* ^Δ*5bp/*Δ*5bp*^. (B) Wild-type and truncated Irx5 proteins produced in the *Irx5* ^Δ*5bp*^ mouse. Both HD and Iro domains are removed in the *Irx5* ^Δ*5bp*^ protein. (C) Immunohistochemistry shows strong reduction in Irx5 expression in *Irx5* ^Δ*5bp/*Δ*5bp*^ spinal cords at e12.5. (D) Double immunohistochemistry for Hb9 and Lhx3 in control and *Irx5* ^Δ*5bp/*Δ*5bp*^ mice at e12.5. (E) Double immunohistochemistry for Foxp1 and Pea3 in control and *Irx5* ^Δ*5bp/*Δ*5bp*^ mice at e12.5. (F-H) Quantification of Hb9+, Foxp1+, or Pea3+ MNs in *Irx5* ^Δ*5bp/*Δ*5bp*^ mice at e12.5 (N=4). * p < 0.05, ** p < 0.01, *** p < 0.001, N. S.: Not significant. (I-J) MN axons labeled with Hb9::GFP in control and *Irx5* ^Δ*5bp/*Δ*5bp*^ mice at e12.5 (N=4). Projections to forelimb (H) and cutaneous maximus (I) muscle.

### *Irx6* limits the number of brachial MNs in the mouse spinal cord

Due to its restricted expression in a subpopulation of LMC neurons (Pea3 pool) (**Fig. 2M-O**), we mutated *Irx6* to test whether it is required for LMC development. We used CRISPR/Cas9 to generate *Irx6* mutant mice that carry a 19bp-long deletion in the 2^nd^ of the six *Irx6* exons (**Fig. 5A**) (see Materials and Methods). Homozygous *Irx6 ^119bp^*^/^*^119bp^* mice are viable. The deletion is predicted to generate a truncated protein lacking both the HD and IRO domains (**Fig. 5B, Supplementary File 1**). Next, we quantified the number of LMC neurons (Foxp1+ cells) at consecutive sections along the rostrocaudal axis of the e12.5 brachial spinal cord. Compared to control *Irx6^+/+^*mice, we found a significant increase in the number of Foxp1+ MNs in rostral sections of the *Irx6 ^119bp^*^/^*^119bp^* spinal cord (**Fig. 5C-D**). We therefore conclude that *Irx6* has a relatively minor role in the developing spinal cord, which is to limit the number of LMC neurons in brachial (rostral) domains.

**Figure 5.**
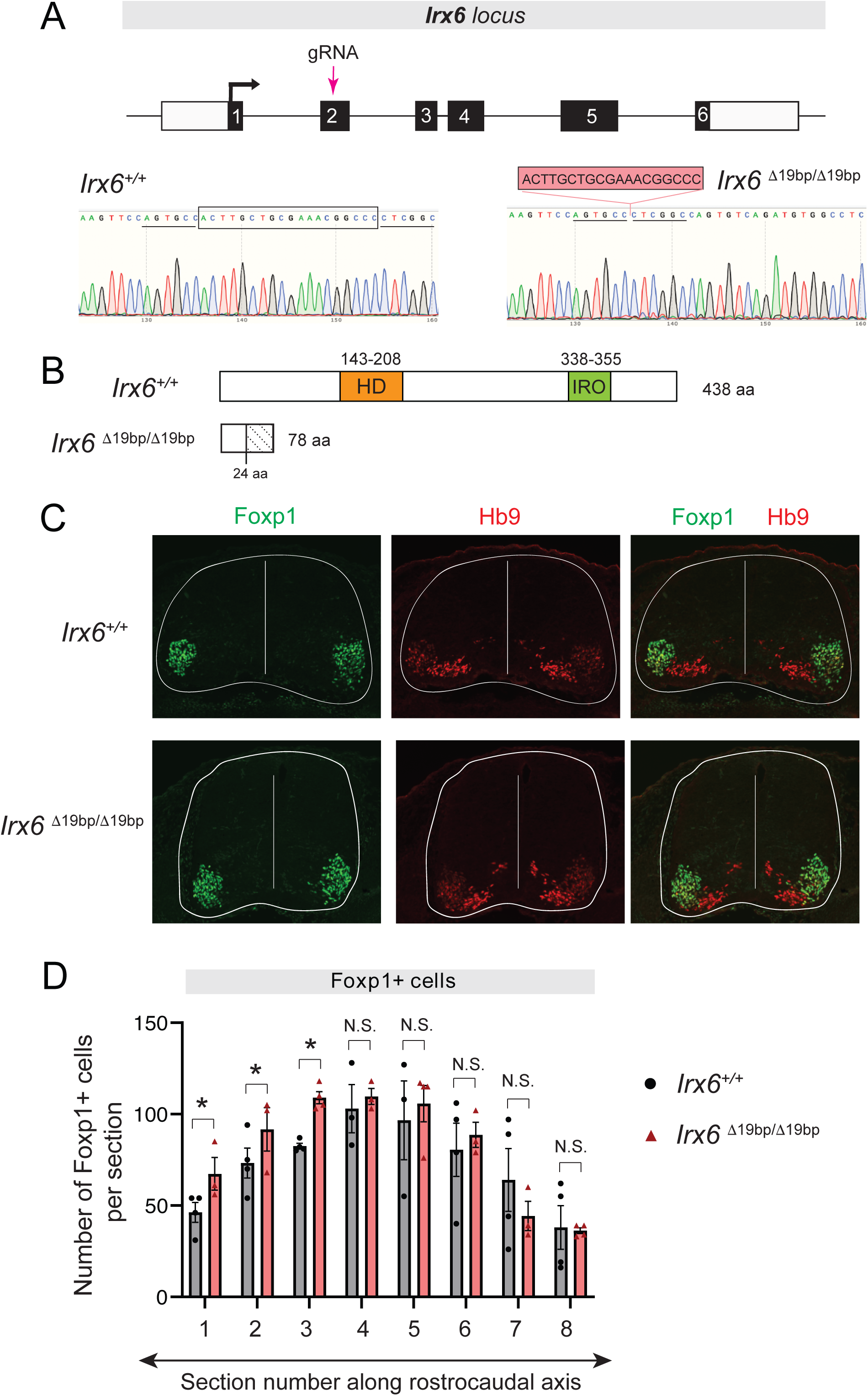
Generation and characterization of *Irx6* mutant mice. (A) A gRNA targets exon 2 of *Irx6* locus. Representative chromatograms of genotyped mice illustrate the 19 bp deletion in the *Irx6* ^Δ*19bp/*Δ*19bp*^ mouse. (B) Wildtype and truncated Irx6 proteins produced in the *Irx6*^Δ*19bp*^ mouse. HD and Iro domains are removed in the *Irx6* ^Δ*19bp*^ protein. (C) Antibody staining for Foxp1-expressing MNs (green) and Hb9-positive MNs (red) in control (*Irx6* ^+/+^) and *Irx6* ^Δ*19bp/*Δ*19bp*^ spinal cords. (D) Quantifications of Foxp1+ MNs along the rostro-caudal axis reveal an increased number of LMC MNs in rostral sections of the *Irx6* ^Δ*19bp/*Δ*19bp*^ spinal cord (N=4). * p < 0.05, ** p < 0.01, *** p < 0.001, N. S.: Not significant.

### Hoxc8 controls *Irx2, Irx5,* and *Irx6* expression in brachial motor neurons

In the developing spinal cord, expression of *Irx3* in progenitor cells is regulated by Shh signaling^30,48^. In post-mitotic neurons, however, little is known about *Irx* gene regulation. At the brachial domain of the mouse spinal cord, the transcription factor *Hoxc8* controls various facets of MN development including axon guidance and terminal differentiation^56,64^. We therefore hypothesized that Hoxc8 may also control *Irx* gene expression in brachial MNs.

To test this, we crossed homozygous mice carrying a conditional *Hoxc8* allele (*Hoxc8^fl/fl^*) to *Olig2^Cre^* mice that enable *Cre* recombinase expression specifically in MN progenitors ^65^, thus effectively removing *Hoxc8* gene activity from their descendant post-mitotic MNs before e12.5^64^. Post-mitotic MNs in mice are generated between e9 - e11 ^66^. Because e12.5 is an early stage of MN differentiation we will refer to the *Olig2^Cre^::Hoxc8 ^fl/fl^* mice as *Hoxc8 MNΔ ^early^*. In the LMC neurons of these mice at e12.5, we found through RNA ISH that expression of *Irx2* is reduced, whereas *Irx5* and *Irx6* expression is undetectable (**Fig. 6A**). These effects are not an indirect consequence of MN cell loss as the total number of brachial MNs is unaffected in *Hoxc8 MNΔ ^early^* animals at e12.5^64^. We conclude that Hoxc8 is required during development (at e12.5 or earlier) to activate *Irx2, Irx5* and *Irx6* expression in young post-mitotic LMC neurons.

**Figure 6.**
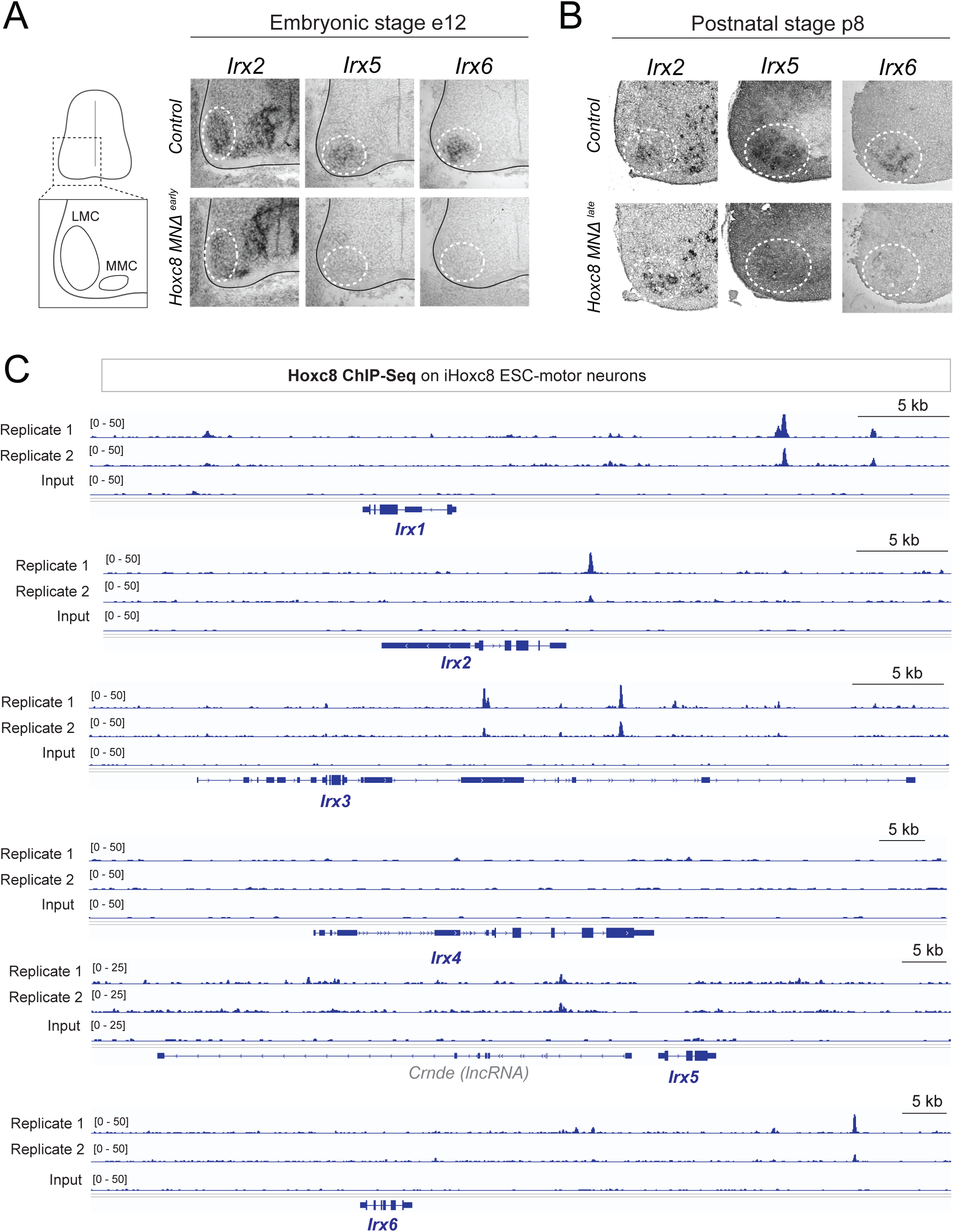
Hoxc8 controls *Irx* expression in spinal motor neurons. (A) RNA ISH shows reduced *Irx2, Irx5* and *Irx6* expression in e12.5 *Hoxc8 MNΔ^early^* spinal cords. (B) At p8, *Irx5* and *Irx6* expression is reduced in *Hoxc8 MNΔ^late^* spinal cords, but *Irx2* expression is unaffected (N=4). (C) Analysis of ChIP-Seq data from iHoxc8 MNs shows Hoxc8 directly binds to the *cis*-regulatory region of *Irx* genes. GEO accession numbers: Input (GSM4226461) and iHoxc8 replicates (GSM4226436, GSM4226437). Snapshots of each gene locus were generated with Integrative Genomics Viewer (IGV, Broad Institute).

How is *Irx* gene expression maintained in post-mitotic MNs? Because *Hoxc8* and *Irx* genes continue to be expressed in LMC neurons during developmental and postnatal stages (**Fig. 2**) ^64,67^, we asked whether Hoxc8 is required to maintain *Irx* expression in LMC neurons. We crossed the *Hoxc8^fl/fl^*mice with the *Chat^IRESCre^* mouse line, which enables efficient gene inactivation in post-mitotic MNs around e13.5 – e14.5 ^64,68^. We refer to the *Chat^IRESCre^::Hoxc8^fl/fl^* animals as *Hoxc8 MNΔ ^late^* because *Hoxc8* depletion in MNs occurs later compared to *Hoxc8 MNΔ ^early^* mice. Interestingly, *Irx2* expression is unaffected in MNs of *Hoxc8 MNΔ ^late^* mice at postnatal day 8 (p8) (**Fig. 6B**). However, expression of *Irx5* and *Irx6* is reduced at p8 (**Fig. 6B**). Altogether, our findings on *Hoxc8 MNΔ ^early^* and *Hoxc8 MNΔ ^late^* mice demonstrate that, in brachial LMC neurons, Hoxc8 is required to induce *Irx2*, *Irx5* and *Irx6* during early development, as well as to maintain *Irx5* and *Irx6* during later developmental stages.

### Hoxc8 acts directly to control *Irx* gene expression in mouse motor neurons

To gain mechanistic insights on *Irx* gene regulation, we analyzed available chromatin immunoprecipitation-sequencing (ChIP-seq) datasets on MNs derived from mouse embryonic stem cells (mESCs), in which *Hoxc8* expression was induced with doxycycline ^69^. In this context, ChIP-Seq for Hoxc8 revealed binding in the *cis*-regulatory region of *Irx1, Irx2, Irx3, Irx5,* and *Irx6* genes. No binding was observed *on Irx4*, the only *Irx* gene not expressed in spinal MNs (**Fig. 1C, 6C**). The ChIP-Seq data, together with our *in vivo* findings in *Hoxc8* MN*Δ* ^early^ and *Hoxc8* MN*Δ* ^late^ mice, strongly suggest that Hoxc8 acts directly to activate *Irx* gene expression in spinal MNs (**Fig. 7E).**

**Figure 7.**
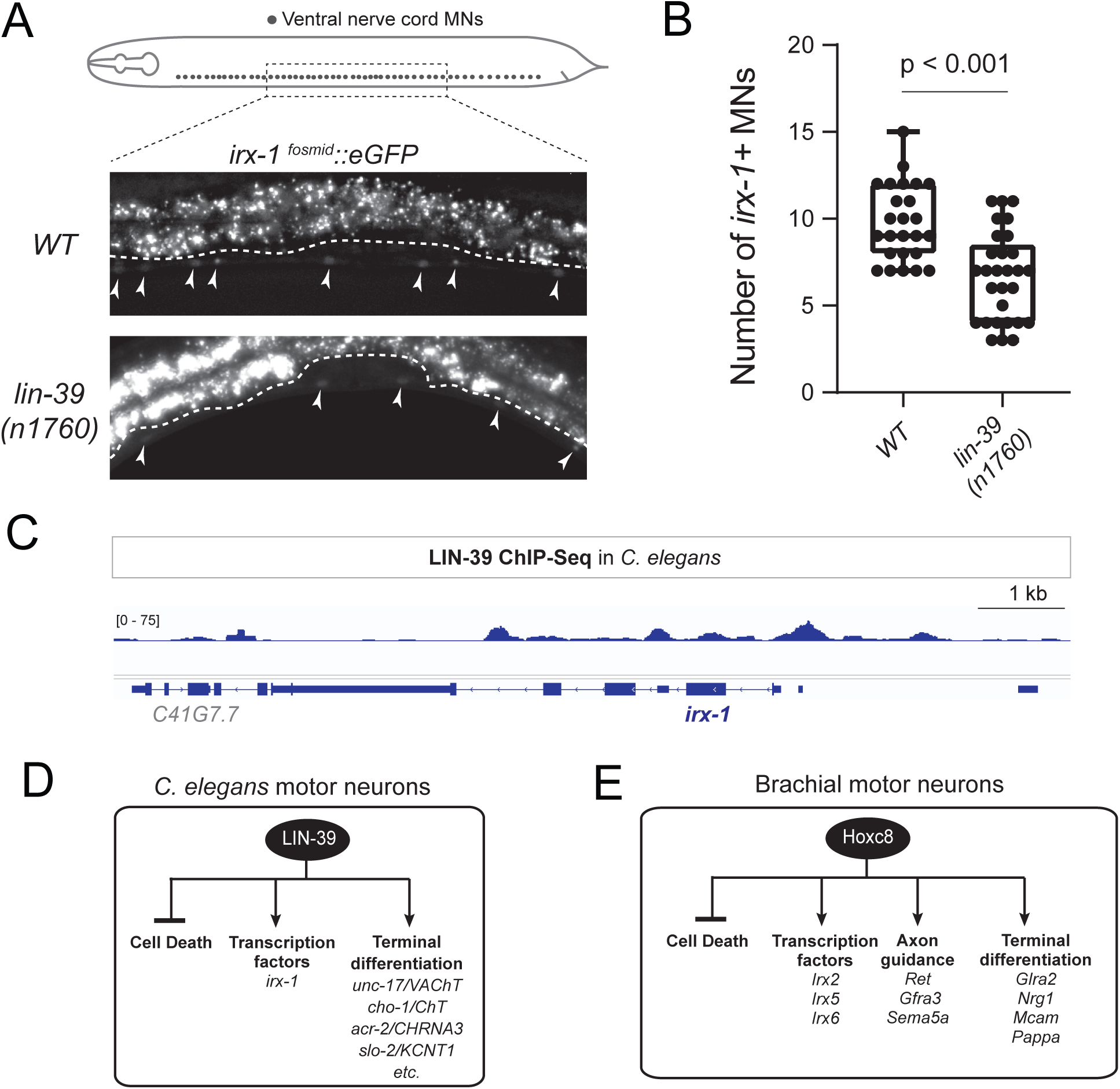
LIN-39/Hox controls *irx-1* expression in *C. elegans* motor neurons. (A) MN cell body position along the *C. elegans* ventral nerve cord. Expression of an *irx-1::eGFP* reporter *(wgIs536*) is decreased in MNs of *C. elegans* animals carrying the *lin-39* (*n1760*) LOF allele. A 300µm region of the ventral nerve cord (VNC) was analyzed at the fourth larval stage (L4). Anterior is left, dorsal is up. Arrowheads point to MN nuclei. Green fluorescence signal is shown in white for better contrast. (B) Quantification of total number of MNs expressing *irx-1* in WT and *lin-39* null mutants. Box and whiskers plot show the min, max and quartiles with single data points annotated. N > 23. (C) ChIP-Seq tracks are shown for LIN-39 on *irx-1*. ChIP-Seq data for LIN-39 come from the modENCODE project^73^. (D) Schematic of known LIN-39 functions and target genes in *C. elegans* MNs. See Discussion for details. A complete list of terminal differentiation genes regulated by LIN-39 in *C. elegans* MNs can be found in ^90,91^. (E) Schematic of known Hoxc8 functions and target genes in brachial MNs. See Discussion for details. Target genes of Hoxc8 with roles in axon guidance and terminal differentiation have been previously described^56,64^.

### LIN-39/Hox directly activates *irx-1* in *C. elegans* motor neurons

Both vertebrate and invertebrate *Irx/Iro* genes are regulated by Hedgehog and TGF-b signaling pathways ^21,48,70^, suggesting control mechanisms regulating *Irx/Iro* gene expression are under strong evolutionary pressure and thereby conserved across distant species. To test whether the role of Hox genes in regulating *Irx/Iro* expression is deeply conserved, we turned to the nematode *C. elegans*, which is evolutionarily separated from mice by hundreds of millions of years (**Fig.1A**). In the *C. elegans* ventral nerve cord (analogous to the mouse spinal cord), MNs that control locomotion co-express the Hox gene *lin-39 (Scr/Dfd/Hox3-5)* and the sole *Irx/Iro C. elegans* ortholog, *irx-1* ^71^. We observed a significant reduction in the number of MNs expressing an *irx-1::gfp* reporter in *lin-39/Hox* null mutant animals at the fourth larval stage (L4) (**Fig. 7A-B**). Because nerve cord MNs that control locomotion are normally generated in *lin-39* mutant animals^72^, we conclude that *irx-1* expression in *C. elegans* MNs requires positive LIN-39/Hox input. Further, we analyzed available LIN-39 ChIP-Seq data at L3^73^, a stage by which all *irx-1*-expressing MNs have been generated. Similar to our Hoxc8 observations in mouse MNs, LIN-39 *(Scr/Dfd/Hox3-5)* binds directly to the *cis*-regulatory region of *irx-1* (**Fig. 7C**). Altogether, our findings in *C. elegans* and mouse MNs uncover an ancient role for Hox proteins in the regulation of *Irx* gene expression (**Fig. 7D-E**).

## DISCUSSION

The expression, function, and regulation of *Irx/Iro* homeobox genes in the nervous system is poorly defined. In this study, we provide a detailed expression map of all six mouse *Irx* genes in the developing spinal cord, revealing unique and overlapping sites of expression in post-mitotic MNs. Further, we uncover a requirement for mouse *Irx2* and *Irx6* in MN development. Last, we show – both in *C. elegans* and mouse MNs – that HOX transcription factors positively regulate *Irx* gene expression, exposing an ancient gene regulatory relationship between two highly conserved families of clustered transcription factors.

### *Irx* genes: a new example of homeobox genes critical for MN development

Locomotion is an ancient behavior displayed by invertebrate and vertebrate animals. It is therefore likely that evolutionary pressure led to a considerable degree of conservation in the molecular mechanisms that control the development and function of various cell types essential for locomotion. This is particularly evident at the level of MNs. Accumulating evidence suggests that highly conserved gene regulatory programs control MN development. For example, the homeobox transcription factor HB9/Mnx1, which is transiently expressed in developing MNs, specifies MN subtype identity in worms, flies, simple chordates, and vertebrates^74–78^. Perhaps the most striking example of conserved regulators of MN development comes from Hox genes, a family of chromosomally clustered homeobox genes. During worm, fly and mouse MN development, region-specific expression of Hox proteins along the anterior-posterior axis of the nervous system is required for the early steps of MN development (e.g., cell specification, circuit assembly)^55^. Here, we show that members of the *Irx* homeobox gene family are expressed in worm, mouse and human MNs, consistent with previous *C. elegans* studies and mouse transcriptomic datasets^67,71,80^. Further, *Irx* genes are expressed in MNs of the little skate, *Leucoraja erinacea*, a marine species of jawed vertebrates, sharing a common ancestor with tetrapods^81^. The remarkable conservation in *Irx* MN expression suggests critical functions. Consistent with this idea, *irx-1* controls MN connectivity in *C. elegans* ^71^, and our data uncover essential roles for *Irx2* and *Irx6* in mouse spinal MN development. Altogether, our findings on *Irx* genes together with previous studies on HB9/Mnx1 and Hox genes lend support to the notion that homeobox genes from different families (e.g., HOX, IRX) play crucial roles in MN development.

### *Irx* genes in the developing spinal cord: Essential and redundant roles

To date, our understanding of *Irx* gene function in the developing spinal cord has been limited to *Irx3*. Distinct classes of spinal neurons are generated at defined D-V positions in response to antiparallel morphogenetic gradients (e.g., Shh, BMP, Wnt)^82^. A set of homeodomain transcription factors is known to act as intermediaries of Shh activity and fall into two classes: class I proteins are repressed by Shh, and class II proteins are activated by it. The expression of these proteins defines distinct progenitor domains in the chick, mouse and human spinal cord^30,48,83^, thereby generating distinct classes of neurons along the D-V axis of the spinal cord. Sharp boundaries between these progenitor domains are maintained via mutual cross-repression of class I and II proteins. In this context, *Irx3* is a class I protein that represses two class II proteins (Olig2, Nkx2.2)^28,30^.

In this study, we describe essential but opposing roles for *Irx2* and *Irx6* genes in spinal MN development. The mouse mutant phenotypes suggest that *Irx2* promotes the generation of LMC (Foxp1+) neurons in the entire brachial domain of the spinal cord (**Fig. 3**), whereas *Irx6* limits the generation of these neurons specifically in rostral domains of the brachial spinal cord (**Fig. 5**). We did not observe any phenotypes in our *Irx5* mutant animals, possibly due to genetic redundancy with *Irx6.* Of note, *Irx5* is known to act redundantly with other *Irx* genes in the context of heart development ^84^. However, the neighboring position of *Irx5* and *Irx6* on the same chromosome prevented us from generating double *Irx5^15bp/15bp^; Irx6^119bp/119bp^* mutant mice by genetic crossing (**Fig. 1B**).

### The *Irx* genes constitute ancient Hox targets in motor neurons

An interesting aspect of *Irx* genes is their conserved patterns of expression. Within vertebrate lineages, the expression of *Irx* orthologs is largely equivalent, suggesting conservation of their *cis*-regulatory elements. For example, all vertebrate *Irx3* orthologs are expressed in equivalent regions of the neural tube and lateral mesoderm ^19,52,85^. Further, both vertebrate and invertebrate *Iro/Irx* genes are regulated by Hedgehog and TGF-beta signaling ^21,48,70^, suggesting that *cis*-regulatory elements controlling the ancestral *Iro/Irx* gene have been evolutionarily conserved. Our findings support this idea, and expand our understating of *Iro/Irx* gene regulation in the following ways.

First, we show that *Irx* genes are regulated by Hox transcription factors both in *C. elegans* and mouse MNs, exposing an ancient gene regulatory relationship. Second, ChIP-Seq data for *C. elegans* LIN-39/Hox and mouse Hoxc8 provide mechanistic insights supporting a model of direct regulation of *Irx* genes by Hox transcription factors. (**Fig. 6-7).**

The selective binding of Hoxc8 to *cis*-regulatory regions of *Irx* genes expressed in MNs (*Irx1, Irx2, Irx3, Irx5, Irx6*) suggests the presence of MN-specific enhancers driving *Irx* expression. The binding regions of LIN-39/Hox and Hoxc8 onto *Irx* genes provide a starting point for future functional assays to identify such enhancers in *C. elegans* and mice. Such assays can be of biomedical relevance. For example, enhancer activity of *Irx3* and *Irx5* in the hypothalamus, a brain region involved in feeding behavior and metabolism, has been linked to obesity ^41–44^. Last, while it is known that Shh represses *Irx3* along the D-V axis, how *Irx* genes are regulated along the anteiro-posterior axis of the spinal cord remains unknown. We found that *Irx* genes are expressed in MNs of the brachial, thoracic, and lumbar domains.

Similar to the action of Hoxc8 in brachial MNs, other Hox proteins may regulate *Irx* gene expression in thoracic and lumbar domain MNs.

### Expanding the repertoire of Hox target genes in the nervous system

HOX transcription factors play fundamental roles in hindbrain and spinal cord development ^55,86,87^. Yet, their downstream target genes in the nervous system remain poorly defined. Our study expands the known repertoire of Hox target genes in the nervous system. Both in *C. elegans* and mouse MNs, Hox transcription factors activate *Irx* gene expression. Intriguingly, the functional roles for Hox in *C. elegans* and mouse MN development appear highly conserved. In *C. elegans*, LIN-39/Hox prevents cell death of a specific MN subtype^88,89^, but is also required more broadly for the terminal differentiation program of ventral nerve cord MNs (**Fig. 7D**) ^90–92^. In the mouse spinal cord, global *Hoxc8* KO studies established that *Hoxc8* is needed to prevent MN cell death ^93^, and conditional KO approaches showed that it promotes axon guidance and terminal differentiation (**Fig. 7E**) ^56,64^. Since global inactivation of either *Hoxc8* or *Irx2* results in decreased numbers of brachial MNs, we speculate that *Irx2* may act as an intermediary factor – downstream of Hoxc8 – to prevent MN cell death (**Fig. 7E**).

### Limitations

We found five *Irx* genes (*Irx1, Irx2, Irx3, Irx5, Irx6*) to be expressed in post-mitotic spinal MNs in the mouse embryo. Although we uncovered a requirement for *Irx2* and *Irx6* in MN development, conditional mutagenesis is needed to determine whether *Irx2* and *Irx6* act at the level of progenitor cells or post-mitotic MNs. Due to their overlapping expression in spinal MNs and reported *Irx* heterodimerization in cardiomyocytes (i.e., Irx3, Irx4, and Irx5 physically interact)^94^, combined mutant analysis is needed to uncover possible redundant roles for *Irx* genes in the spinal cord. Last, we recently reported continuous *Irx* gene expression in mouse spinal MNs during early postnatal and adult life^64,67^. Hence, the *Irx* expression map we describe here is likely preserved throughout post-embryonic life. Inducible mutagenesis is therefore needed to determine the complete repertoire of *Irx* gene functions in MNs at different stages of embryonic and postnatal life.

## ACKNOWLEDGEMENTS

We thank members of the Kratsios lab (Ian Weigle, Mira Antonopoulos, Jayson J. Smith, Nidhi Sharma, Filipe Marques, Honorine Destain, Manasa Prahlad) for comments on the manuscript and Dr. Jeremy Dasen (NYU) for providing antibodies (rabbit anti-Foxp1, rabbit anti-Lhx3, rabbit anti-Hb9, rabbit anti-Isl1/2). We thank Jayson J. Smith for help with figure design and the University of Chicago Transgenic Mouse Facility (RRID:SCR_019171) and its technical director (Linda Degenstein) for generating *Irx2, Irx5* and *Irx6* mutant mice using CRISPR/Cas9 genome editing. The *C. elegans* strains were provided by the CGC, which is funded by NIH Office of Research Infrastructure Programs (P40 OD010440). This work was supported by the Lohengrin Foundation and the National Institute of Neurological Disorders and Stroke (NINDS) (Award Number: R01NS116365 to P.K.).

## AUTHOR CONTRIBUTIONS

Conceptualization: C.C., P.K.; Methodology: C.C., S.A., Y.C., W.F., K.T.; Investigation: C.C., S. A., Y.C., W.F., K.T.; Formal analysis: C.C., S.A., Y.C., W.F., K.T.; Visualization: C.C., P.K.; Funding acquisition: P.K.; Writing original draft: P.K.; Writing - review and editing: C.C., S.A., Y.C., W.F., K.T.; Supervision: P.K.

## DATA ACCESSIBILITY

All data generated or analyzed in this study are included in the manuscript and supporting files.

## MATERIALS AND METHODS

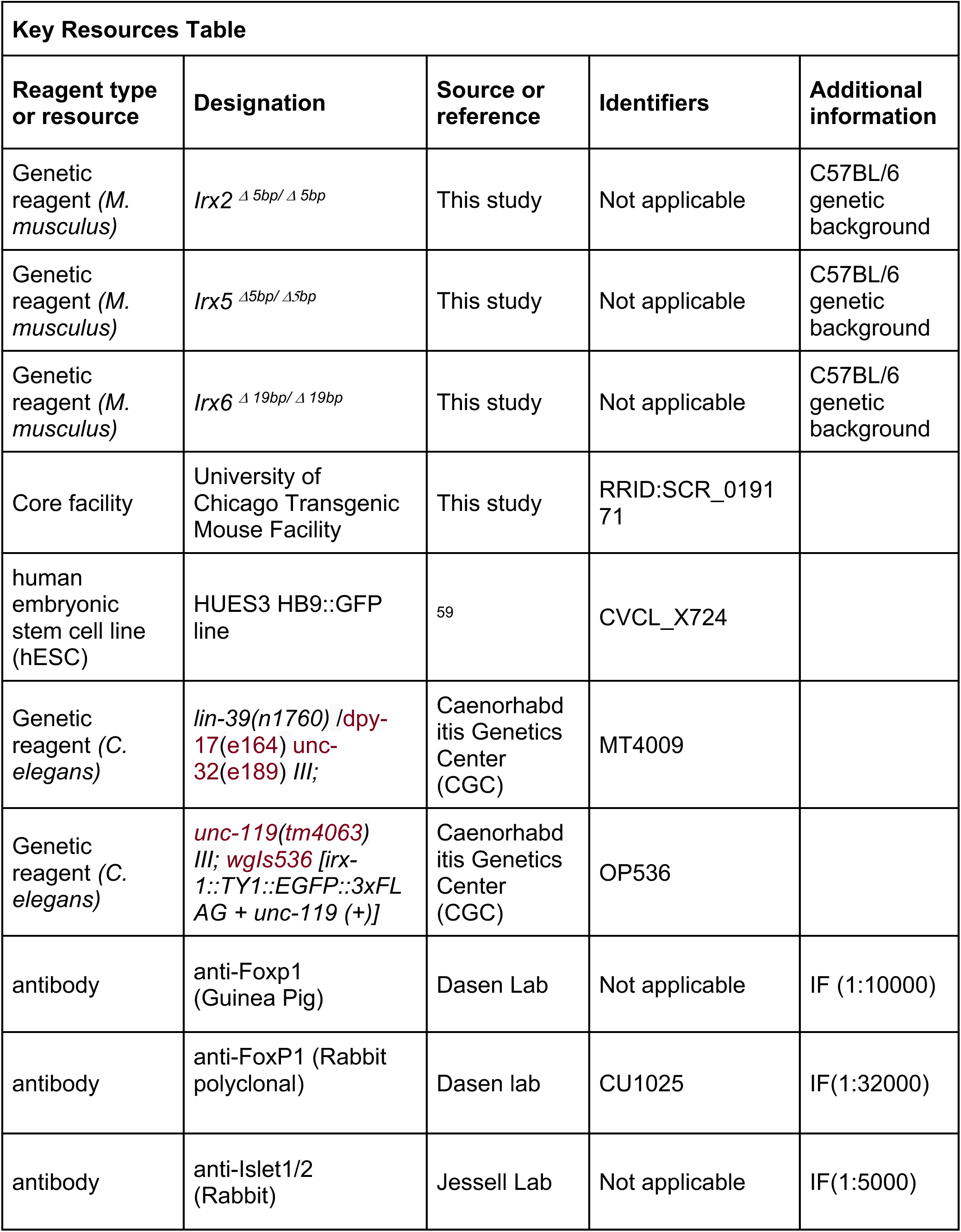
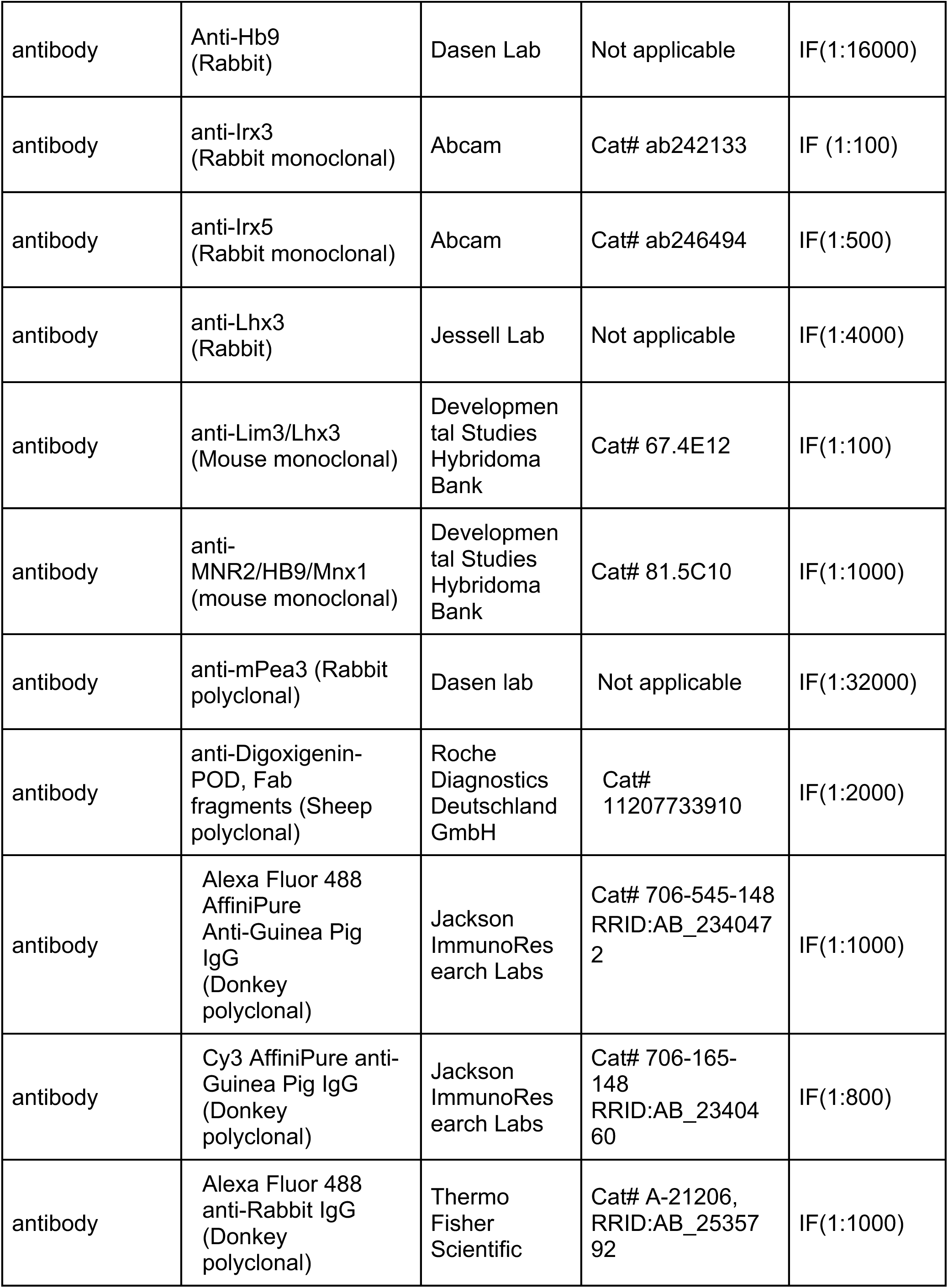
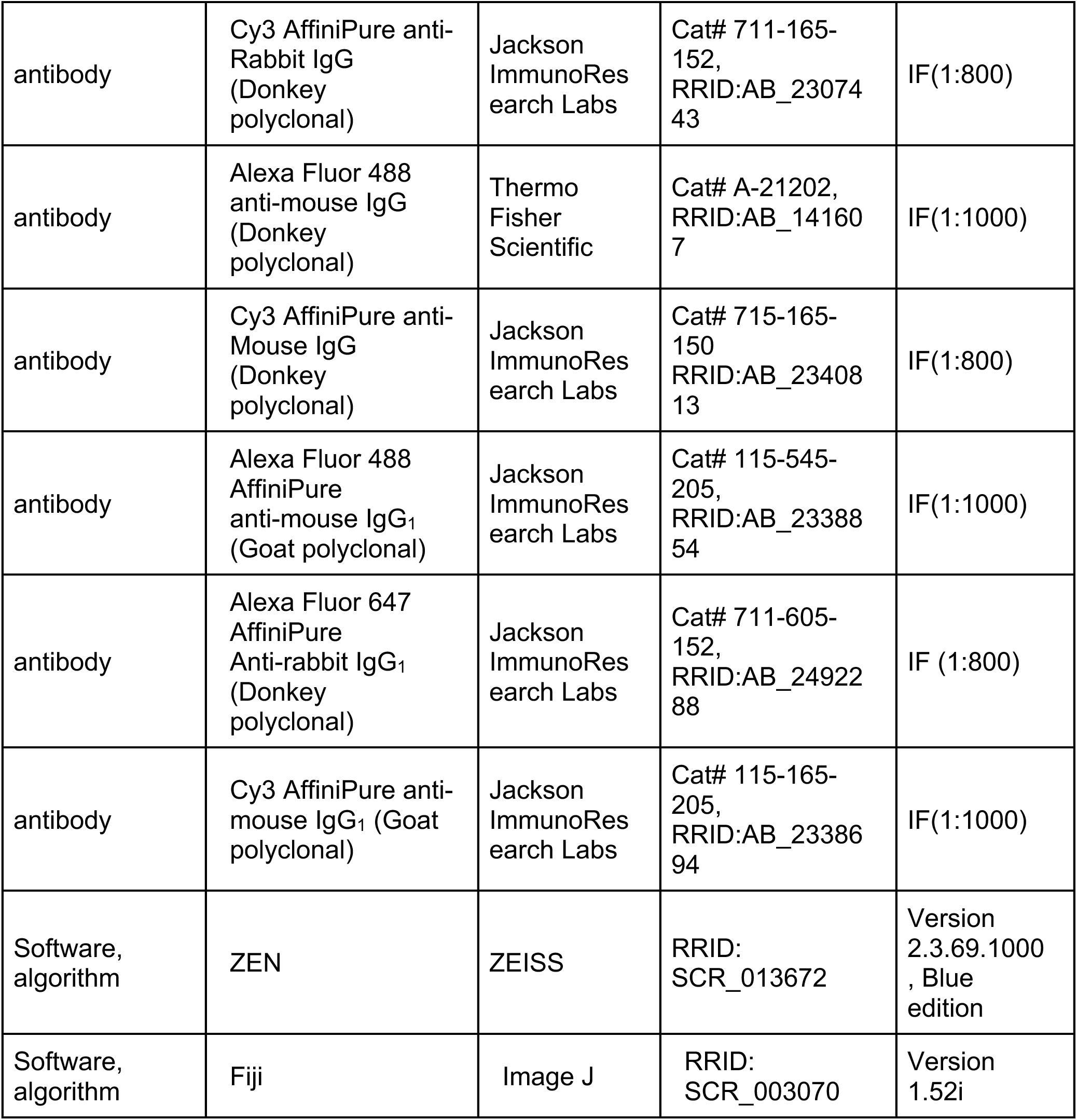

### Mouse husbandry and genetics

All mouse procedures were approved by the Institutional Animal Care and Use Committee (IACUC) of the University of Chicago (Protocol # 72463). Morning (9AM) of the day on which a vaginal plug was seen was termed embryonic day 0.5 (E0.5).

The *Irx2* mutant mice were generated at the Transgenics/ES Cell Technology Mouse Core Facility of the University of Chicago. A gRNA targeting exon 2 of *Irx2* (gRNA: GTACCGTAAGAACGCTACGC) was injected into mouse zygotes together with a Cas9- expressing plasmid. Founder animals were genotyped by sequencing and a 5 bp deletion was detected in exon 2 of the *Irx2* locus (*Irx2 ^15bp/15bp^*). In the F1 generation, heterozygous *Irx2 ^15bp/15bp^* mice were used for crosses to establish a homozygous mutant line. Primers for genotyping the *Irx2 ^15bp/15bp^* allele: Forward primer 5’-ACAGGCTGTTGTGGGTTCC-3’, Reverse primer 5’-CATCCTGTGCCTTGTCTGAA-3’. Using similar CRISPR/Cas9 procedures, we generated the *Irx5 ^15bp/15bp^* and *Irx6 ^119bp/119bp^* mice. gRNA targeting *Irx5:* 5’-GTACCAGCCGTCCGCCTCGC-3’ Primers for genotyping the *Irx5 ^15bp/15bp^* allele:

Irx5 Forward 5’-AGAAGCCAGGTGCCCTCTC-3’ Irx5 Reverse 5’-CAGCTCACACTCACCACGTAA-3’ *gRNA targeting Irx6:* 5’-CTTGCTGCGAAACGGCCCCT-3’ Primers for genotyping the *Irx6^119bp/119bp^* allele: Irx6 Forward 5’-GAGATTCTGCACCTGGTGGT-3’ Irx6 Reverse 5’-CTTGTCCCCAGACAAGGTCACAG-3’

### Mouse tissue collection, processing, and immunofluorescence

Mouse embryos were harvested at E12.5 and fixed for 1.5 hr in 4% paraformaldehyde in PBS at 4C. Embryos were then washed in ice-cold PBS, equilibrated overnight in 30% sucrose/PBS at 4C, embedded in optimal cutting temperature (OCT) compound, and sectioned at 12um on a Leica CM3050 S cryostat. For immunofluorescence staining, sections were briefly washed in PBS, blocked for 30 min in 1% BSA in PBST (PBS with 0.1% Triton X-100) at room temperature, then incubated overnight in primary antibodies (diluted in 0.1% BSA/PBT) at 4°C. Sections were then washed in PBST, incubated in secondary antibodies in PBST for 1 hr at room temperature, and washed again in PBST before applying the Prolong Diamond antifade mounting medium (Invitrogen Cat #: P36971) and a coverslip. Primary and secondary antibodies are listed in Key Resources Table.

### RNA *in situ* hybridization

E12.5 embryos and p8 spinal cords were fixed in 4% paraformaldehyde for 1.5–2 h and overnight, respectively, placed in 30% sucrose/PBS overnight (4 °C), and embedded in OCT compound. Cryosections were generated and processed for *in situ* hybridization or immunohistochemistry as previously described ^95,96^.

### RNA fluorescent *in situ* hybridization coupled with antibody staining

Cryosections were postfixed in 4% paraformaldehyde and washed in PBS. Endogenous peroxidase was blocked with a 1% H_2_O_2_ solution and cryosections were permeabilized in PBS/0.1% Triton-X100. Next, sections were hybridized with a DIG-labeled RNA probe overnight at 72 °C, and washed in SSC. Then, the anti-DIG antibody conjugated with peroxidase (Roche) and primary antibody against Foxp1 (rabbit anti-Foxp1, Dr. Jeremy Dasen), Irx3 (rabbit anti-Irx3, Abcam), or Lhx3 (mouse anti-Lim3, Developmental Studies Hybridoma Bank) were applied overnight (4 °C) to the sections. The next day, the sections were incubated with the secondary antibody (Alexa 488 donkey anti-rabbit IgG, Life Technologies) or Alexa 488 goat anti-mouse IgG1, Jackson ImmunoResearch Labs) and detection of RNA was performed using a Cy3 Tyramide Amplification system (Perkin Elmer). Images were obtained with a high-power fluorescent microscope (Zeiss Imager V2) and analyzed with Fiji software ^97^.

For double RNA fluorescence *in situ* hybridization, two different probes were used, one labeled with digoxigenin and the other with biotin. Anti-DIG-peroxidase conjugated (Roche) and anti-biotin-peroxidase conjugated antibodies were used (Vector Labs). RNA was detected using a fluorescein and a Cy3 Tyramide Amplificafion system (Perkin Elmer).

### Forelimb grip strength test

The forelimb strength of male and female mice was measured using a grip strength meter from Bioseb (model BIO-GS3) ^98^. We assayed control and *Irx2 ^15bp/15bp^* mice at the age of 6.6 to 7.5 months, following the manufacturer’s protocol. In brief, the meter was positioned horizontally on a heavy metal shelf (provided by the manufacturer), assembled with a grip grid. Mice were held by the tail and lowered towards the apparatus. The mice were allowed to grasp the metal grid only with their forelimbs and were then pulled backwards in the horizontal plane. The maximum force of grip was measured, and we used the average of six measurements for analysis. Force was measured in Newtons and grams. The experimenter was blind to the genotypes.

### Differentiation of human motor neurons from embryonic stem cells

For the directed differentiation of embryonic stem cells (HUES3 HB9::GFP line) to post- mitotic spinal MNs, we followed a small molecule-based protocol as previously described^99^.

To induce the neuralization in the culture, we used the TGF-β inhibitor SB43154 (10uM) and the Bone Morphogenetic Pathway (BMP) inhibitor LDN-193189 (100nM). In parallel, to achieve the appropriate patterning, we included in the medium retinoic acid (RA, 1uM) and the Smoothened-Agonist (SAG, 1uM) to promote caudalization and ventralization, respectively. At the second stage of the protocol (days 6-14) where MN terminal differentiation takes place, we replaced the SB43154 and LDN-193189 molecules with the (indirect) inhibitor of the Notch pathway DAPT (5uM) and the FGFR1 inhibitor SU5402 (4uM) in order to force neuronal progenitors to exit the cell cycle. After day 14, the post-mitotic MNs were replated in the presence of neurotrophic factors (BDNF, GDNF, CNTF; 10ng/ml all) for their final maturation.

### Immunocytochemistry on human motor neurons

Motor neurons derived from the HUES3 HB9::GFP line were plated on day 14 on top of 1.5mm glass coverslips (Neuvitro, Fisher scientific) coated with PDL (0.1mg/ml, 37°C overnight) and Laminin (15ug/ml, 37°C for 3h) at a density of 80,000cells/coverslip. Cells were washed once with PBS and fixed with 4% PFA in PBS for 20 min at RT. Following fixation, cells were washed three times with PBS and permeabilized with PBS-Triton X-100 0.2% for 45min at RT. To block the non-specific binding of primary antibody to unrelated epitopes, cells were treated with 10% normal donkey derum (Jackson ImmunoResearch) in PBS-Triton X-100 0.1%, for 1h at RT and then incubated with primary antibody, diluted in 2% Normal Donkey Serum in PBS-TritonX-100 0.1%, overnight at 4°C. The next day, coverslips were washed three times with PBS and incubated with the fluorophore-conjugated secondary antibody (Alexa Fluor 488, or Alexa Fluor 647, all from Invitrogen or Jackson ImmunoResearch) at a dilution 1:1,000 in 2% Normal Donkey Serum in PBS-Triton X-100 0.1%, for 2h at RT protected from light. Cells were washed once with PBS, incubated with DAPI 1:1,000 in PBS for 10min at RT, then washed again three times with PBS. Coverslips were mounted on glass slides (Fisher Scientific) using Fluoromount-G (Southern Biotech).

### Quantification of *irx-1* reporter gene expression in *C. elegans* MNs

Worms were grown at 20°C on nematode growth media (NGM) plates seeded with bacteria (*E.coli* OP50) as food source ^100^. Animals carrying the *lin-39 (n1760)* mutant allele were crossed with animals carrying the *wgIs536 [irx-1::TY1::EGFP::3xFLAG + unc-119 (+)]* fosmid- based reporter for *irx-1.* Homozygous animals for the *lin-39 (n1760)* allele and the *wgIs536* reporter were anesthetized at the fourth larval stage (L4) using 100mM of sodium azide (NaN_3_) and mounted on a 4% agarose pad on glass slides. Images were taken using an automated fluorescence microscope (Zeiss, Axio Imager Z2). Several z-stack images (each ∼1 µm thick) were acquired with Zeiss Axiocam 503 mono using the ZEN software (Version 2.3.69.1000, Blue edition). Representative images are shown following max-projection of 1-8 µm Z-stacks using the maximum intensity projection type. Image reconstruction was performed using Fiji ^97^.

### Statistical analysis

For data quantification, graphs show values expressed as mean ± standard error of the mean (SEM). All statistical analyses were performed using unpaired *t*-test (two-tailed) (GraphPad Prism software). Differences with *p* < 0.05 were considered significant.

## LEGENDS OF SUPPLEMENTARY FILES

Supplementary Table 1: Summary of Irx antibodies used for this study.

Supplementary file 1: Predicted ORF shifts in *Irx2, Irx5* and *Irx6* mutant mice.

Supplementary file 2: RNA ISH probe sequences used in this study.

---

**Supplementary Figure 1.**
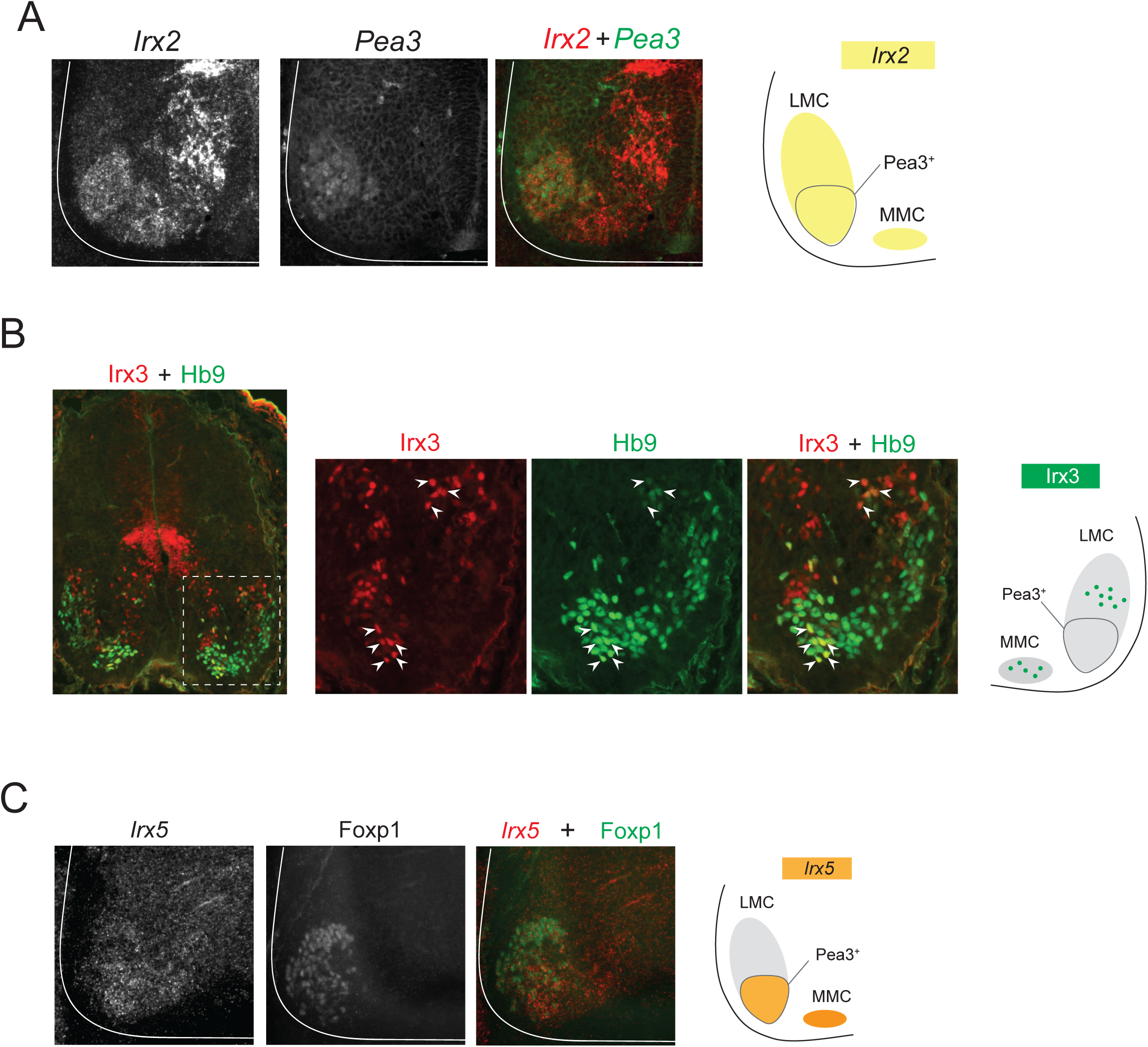
Additional characterization of *Irx2*, Irx3 and *Irx5* expression. (**A-C**) Expression analysis of the brachial spinal domain (C4-T1) in an e12.5 wild-type mouse. (**A**). Two-color RNA FISH analysis shows *Irx2* expression (red) is detected in the ventral LMC at Pea3-expressing MNs (green). Fluorescent signal for *Irx2* (red) and *Pea3* (green) is shown in white for better contrast. (B) Double immunostaining shows Irx3 (red) is detected in small populations of Hb9-expressing MNs (green) of the MMC and dorsal LMC but not in ventral LMC. (C) *Irx5* mRNA (red) colocalizes with Foxp1 protein (green) in ventral LMC domain. (LMC: lateral MC; MMC: Medial MC). Fluorescent signal for *Irx5* (red) and Foxp1 (green) is shown in white for better contrast.

**Supplementary Figure 2.**
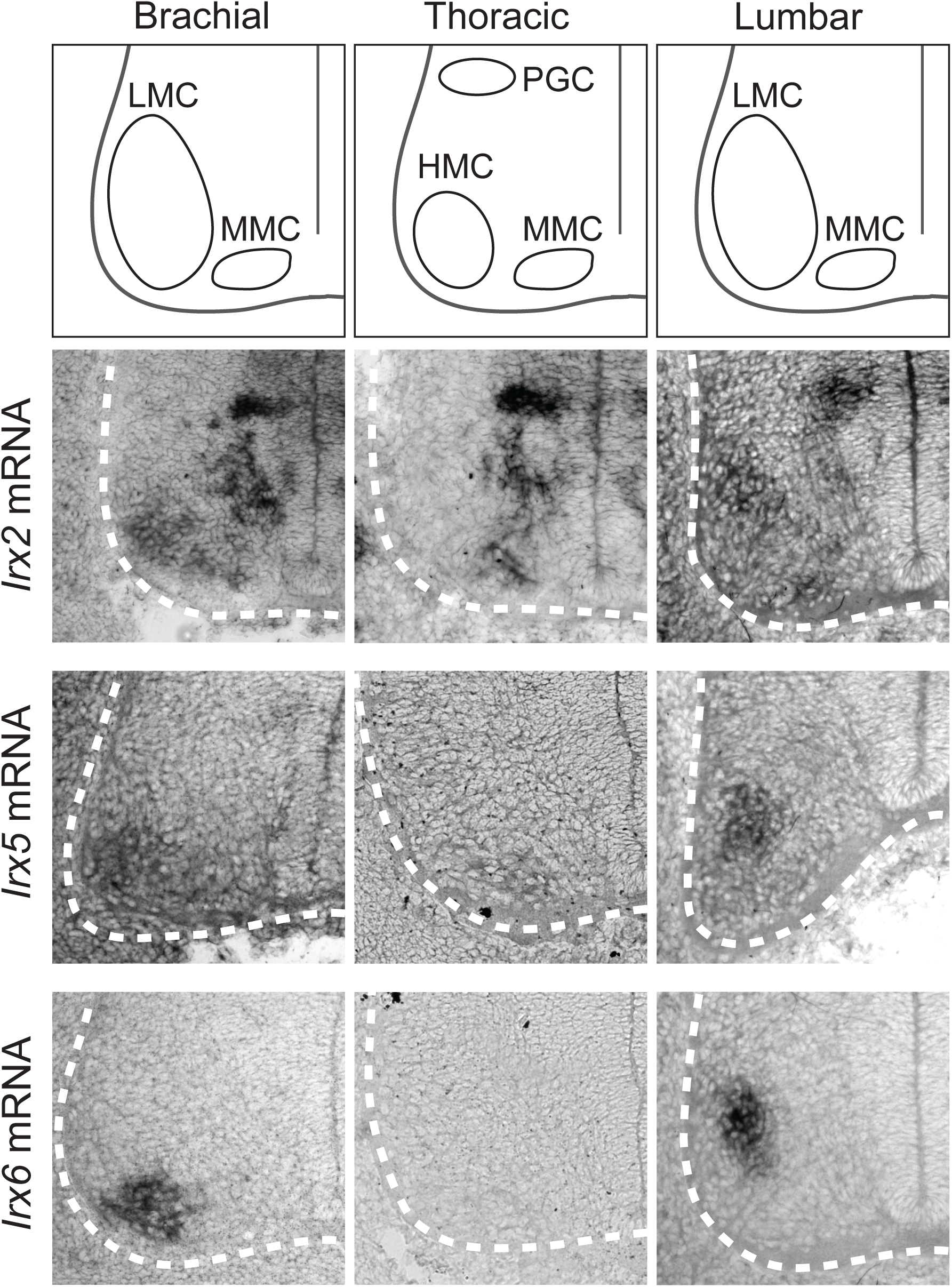
Expression of *Irx2, Irx5*, and *Irx6* at brachial, thoracic and lumbar motor neurons. RNA ISH showing expression of *Irx2* at the brachial, thoracic and lumbar domains of a wild-type e12.5 spinal cord (N = 4). Expression of *Irx5* and *Irx6* is mainly detected at brachial and lumbar regions. *Irx2, Irx5*, and *Irx6* are expressed in the ventrolateral region, which is populated by MNs. A schematic representation of motor column (MC) location is provided at the top (LMC: lateral MC; MMC: Medial MC, HMC: Hypaxial MC; PGC: Preganglionic MC).

**Supplementary Figure 3.**
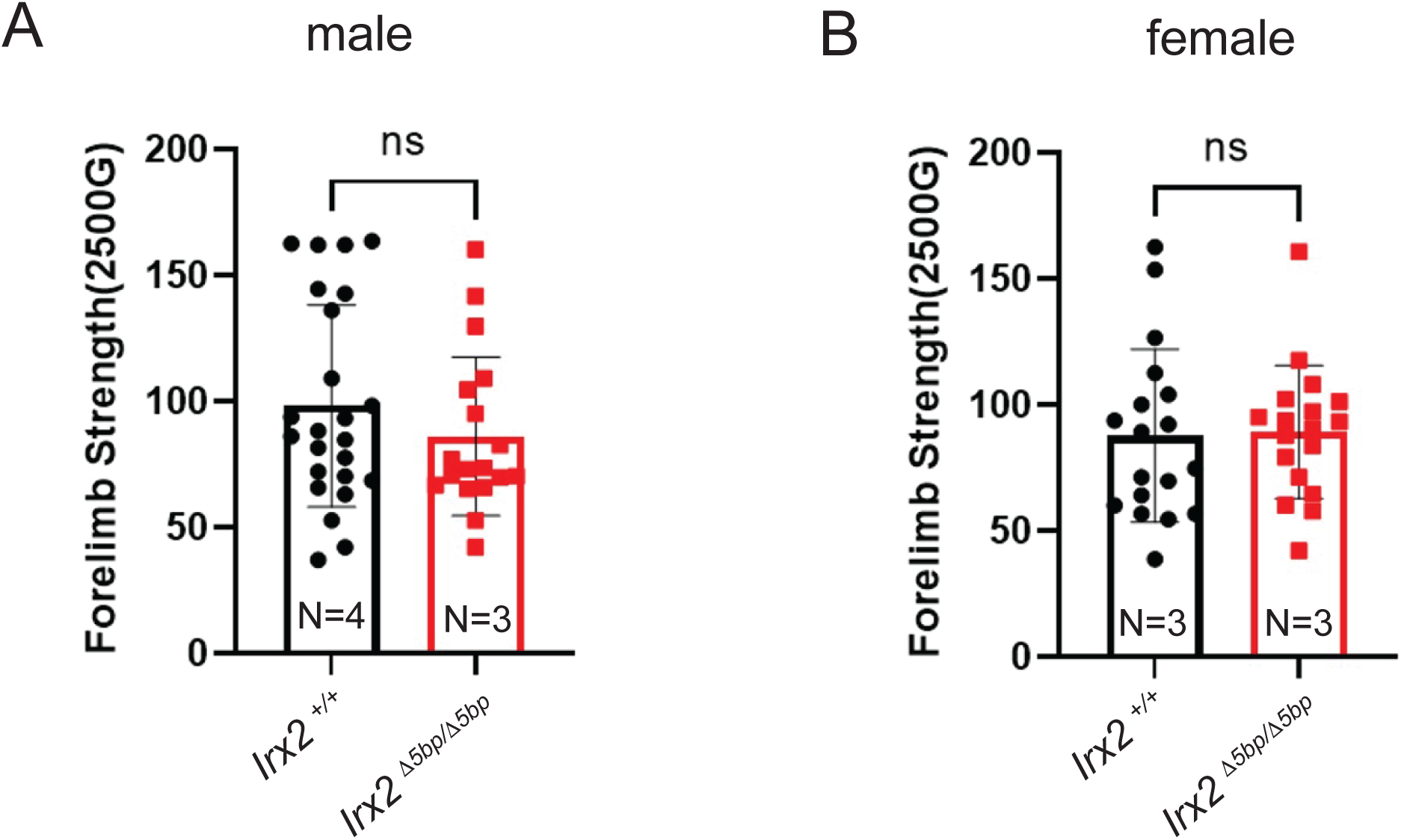
Forelimb grip strength analysis on control and Irx2 ^Δ5bp/Δ5bp^ mice. Control (*Irx2 ^+/+^*) and *Irx2* ^Δ5bp/Δ5bp^ adult mice between 6.5 and 7.5 months old were analyzed. Results are shown for male (A) and female (B) mice. N = number of animals. n.s: Not significant. The maximum force of grip was measured six times for each animal and all values are displayed on the graph. Force was measured in grams (G). The experimenter was blind to the genotypes. See Methods for details.

